# Fluorescent reporters for functional analysis in rice leaves

**DOI:** 10.1101/649160

**Authors:** Leonie H. Luginbuehl, Sherif El-Sharnouby, Na Wang, Julian M. Hibberd

## Abstract

Fluorescent reporters have facilitated non-invasive imaging in multiple plant species and thus allowed analysis of processes ranging from gene expression and protein localization through to cellular patterning. However, in rice, a globally important crop and model species, there are relatively few reports of fluorescent proteins being used in leaves. Fluorescence imaging is particularly difficult in the rice leaf blade, likely due to a high degree of light scattering in this tissue. To address this, we investigated approaches to improve deep imaging in mature rice leaf blades. We found that ClearSee treatment, which has previously been used to visualise fluorescent reporters in whole tissues of plants, led to improved imaging in rice. Removing epidermal and subtending mesophyll cell layers was faster than ClearSee, and also reduced light scattering such that imaging of fluorescent proteins in deeper leaf layers was possible. To expand the range of fluorescent proteins suitable for imaging in rice, we screened twelve whose spectral profiles spanned most of the visible spectrum. This identified five proteins, mTurquoise2, mClover3, mNeonGreen, mKOκ and tdTomato that are robustly expressed and visible in mesophyll cells of stably transformed plants. Using microparticle bombardment, we show that mTurquoise2 and mNeonGreen can be used for simultaneous multicolour imaging of different sub-cellular compartments. Overall, we conclude that mTurquoise2, mClover3, mNeonGreen, mKOκ and tdTomato are suitable for high resolution live imaging of rice leaves, both after transient and stable transformation. Along with the rapid microparticle bombardment method, which allows transient transformation of major cell types in the leaf blade, these fluorescent reporters should greatly facilitate the analysis of gene expression and cell biology in rice.

**One sentence summary:** We report five fluorescent reporters suitable for functional analysis in rice leaves.

## Introduction

The availability of fluorescent proteins has advanced our understanding of biological processes such as membrane trafficking and the subcellular localisation of proteins, the spatial and temporal regulation of gene expression, and the dynamics of cellular patterning and organ development. Unlike other reporters, including β-glucuronidase (GUS) and luciferase, fluorescent proteins allow the visualisation and relative quantification of proteins in live tissue without the need to add a substrate or co-factor (Jefferson et al., 1987; Luehrsen et al., 1992; Haseloff, 1998; Hanson and Köhler, 2001). In addition, the development of a wide range of colour variants based on the original green fluorescent protein (GFP) and the discovery of new fluorescent proteins have enabled multicolour imaging and the simultaneous visualisation of multiple processes in the same tissue, cell, or subcellular compartment (Prasher et al., 1992; Shaner et al., 2005). Fluorescent reporters are also an important component of many experimental tools, such as protein tagging for chromatin immunoprecipitation (De Folter et al., 2007) and fluorescence-activated cell sorting (Birnbaum et al., 2005).

In many plant species, imaging of fluorescent proteins is commonly reported for all major plant organs (Cutler et al., 2000; Bellucci et al., 2005; Mohanty et al., 2009). However, in rice, whilst they are routinely used for analysis in roots and other non-green tissues (Kobae and Hata, 2010; Wu et al., 2016; Chu et al., 2018; Kim et al., 2002; Abiko et al., 2013), there are fewer examples of fluorescent reporters being used in leaves (Dangol et al., 2017; Wu et al., 2016; Meng et al., 2014; Johnson et al., 2005; Reynoso et al., 2018; Wang et al., 2018). Most of these reports show fluorescent protein expression in the leaf sheath (Dangol et al., 2017; Wu et al., 2016; Meng et al., 2014; Wang et al., 2018), but only very few show expression in the leaf blade (Johnson et al., 2005; Reynoso et al., 2018). Importantly, in the leaf blade, expression has only been reported in cells at the leaf surface and not in deeper leaf layers (Johnson et al., 2005; Reynoso et al., 2018). This is likely due to the high degree of light scattering arising from the waxy cuticle of the rice leaf blade, which impedes high resolution fluorescent protein visualisation in deeper layers (Qin et al., 2011; Clark and Lister, 1975). In addition to the difficulties that are associated with deep tissue imaging in rice leaves, a limited number of fluorescent reporters, including GFP, yellow fluorescent protein and red fluorescent protein have been reported to be expressed in rice leaves (Dangol et al., 2017; Reynoso et al., 2018). Thus, available tools for fluorescence imaging in rice leaves are currently limited, hindering the analysis of biological processes such as leaf development, photosynthesis, and chloroplast biogenesis.

The use of fluorescent reporters relies on the ability to stably or transiently transform plant tissue with the required expression vectors. Although rice is widely used as a model due to its relatively small and sequenced genome (Goff et al., 2002; Matsumoto et al., 2005), the availability of a range of genetic resources, and the high level of genomic synteny to other cereals (Bennetzen and Freeling, 1997), stable transformation is labour-intensive and time-consuming (Supartana et al., 2005; Toki et al., 2006; Hiei and Komari, 2008). Thus, transient transformation of rice protoplasts or heterologous expression systems such as *Arabidopsis thaliana* or *Nicotiana benthamiana* are routinely employed to characterize the function of rice proteins (Lam et al., 2007; Kitajima et al., 2009; Zhang et al., 2011). Although less time-consuming, these approaches do not allow protein function to be analysed in cells of intact rice tissues such as fully developed leaves. Transient transformation by microparticle bombardment is an alternative approach that has been shown to enable fluorescence reporter-based analysis in the rice root and leaf sheath but not in the rice leaf blade (Wang et al., 2018).

Here, we tested and optimised tools that enable the expression and imaging of fluorescent proteins in all layers of the rice leaf blade. We tested two methods for deep imaging in rice leaves, clearing using the ClearSee protocol (Kurihara et al., 2015) or scraping the leaf surface, and found that both methods improve visualisation of fluorescent reporters in deeper leaf tissue layers. To expand the range of fluorescent proteins available for imaging in rice, we generated stable transformants expressing twelve reporters with distinct spectral characteristics and tested the extent to which each was visible in leaves. From this analysis we identified five fluorescent proteins that are robustly detectable in rice mesophyll cells. Lastly, microparticle bombardment was optimised to provide a fast and simple method to transiently transform different cell types in the rice leaf blade and other organs of rice. Using fluorescent reporters in stable lines or after transient transformation should facilitate our ability to analyse biological processes such as gene expression, protein subcellular localisation and protein-protein interactions in rice, a globally important crop and model species.

## Materials and Methods

### Cloning and construct design

Fluorescent proteins that were chosen as candidate reporters were selected for analysis primarily based on their relative brightness (Figure S1), but also taking into consideration parameters such as resistance to photobleaching, thermostability, rapid maturation time, and minimal spectral overlap with chlorophyll autofluorescence. Coding sequences of twelve fluorescent proteins selected were obtained from the following sources: mTurquoise2 (Goedhart et al., 2012), mTFP1 [GenBank accession number DQ676819.1, (Ai et al., 2006)], mClover3 (Bajar et al., 2016), mNeonGreen [GenBank accession number KC295282.1, (Shaner et al., 2013)], mCitrine (Griesbeck et al., 2001), mYPet [the monomerizing mutation A206K was introduced into the YPet sequence (Nguyen and Daugherty, 2005) to generate mYPet], mKOκ [7 amino acid mutations (K49E, P70V, K185E, K188E, S192D, S196G, L210Q) were introduced into the mKO sequence with GenBank accession number AB128821.1 to generate mKOκ, (Tsutsui et al., 2008)], tdTomato (Shaner et al., 2004), TagRFP-T [GenBank accession number EU582019.1, (Shaner et al., 2008)], mRuby3 (Bajar et al., 2016), mCardinal (Chu et al., 2014), mKate2 (Shcherbo et al., 2009). Prior to synthesis for GoldenGate cloning (Weber et al., 2011), coding sequences were modified such that the codon encoding valine at position 2 of each open reading frame was mutated to code for an alanine (GCG) to achieve a partial match to the Kozak consensus sequence used in monocots (Nakagawa et al., 2008; Joshi et al., 1997; Gupta et al., 2016), codon-optimized for rice using the codon optimization tool by Integrated DNA Technologies, and domesticated for the GoldenGate cloning system. Where relevant, the GFP cryptic intron (Haseloff et al., 1997) and potential intron sites identified by NetPlantGene Server (Hebsgaard et al., 1996) were neutralized. Each fluorescent protein module was synthesized as an N-terminal tag and fused at the C-terminal end with a GGAGSGAGG amino acid linker to a codon-optimised N7 nuclear localisation signal peptide (Cutler et al., 2000). These modules were assembled with Level 1 modules for the maize *PEPC* promoter (Matsuoka et al., 1994) and *Cauliflower Mosaic Virus 35S* terminator, then combined with the hygromycin resistance gene into the Level 2 binary vector pICSL4723.

For transient expression by microparticle bombardment, the chloroplast targeting sequence *RC2* (Shen et al., 2017) and sequence for the plasma membrane localised protein *OsPIP2.1* (Dangol et al., 2017) were domesticated for the GoldenGate cloning system and synthesised. The binary vector pICSL4723 was assembled through ligation of the *ZmUBIQUITIN* or the *OsACT2* promoter sequences (Cornejo et al., 1993; He et al., 2009); the coding sequences for mTurquoise2 or mNeonGreen; the nuclear, chloroplast, or plasma membrane targeting sequences; and the *NOS* or *CaMV35S* terminator sequences. Assembly of the *GUS* reporter plasmid used for microparticle bombardment has been described previously (Brown et al., 2011).

### Stable transformation of rice

*Oryza sativa spp. japonica* cultivar Kitaake was transformed using *Agrobacterium tumefaciens* as described previously (Hiei and Komari, 2008) with several modifications. Seeds were de-husked and sterilised with 2.5% (v/v) sodium hypochlorite solution for 15 minutes. Seeds were placed on nutrient broth (NB) medium with 2 mg/L 2,4-dichlorophenoxyacetic acid at 30°C in the dark for 3 to 4 weeks to induce callus formation. Actively growing calli were co-incubated with *A. tumefaciens* strain LBA4404 in the dark at 25°C for 3 days. Calli were selected on NB medium containing 35 mg/L hygromycin B for 4 weeks, and proliferating calli regenerated on NB medium containing 10 mg/L hygromycin B for 4 weeks at 28°C under continuous light conditions. Hygromycin resistant plants were screened by PCR with primers amplifying the hygromycin phosphotransferase gene. Positive transformants were transferred to a 1:1 mixture of top soil and sand, and then grown in a greenhouse under natural light conditions but supplemented with minimum light intensity of 390 μmol m^−2^ s^−1^, 60% humidity, temperatures of 28°C and 23°C during the day and night, respectively, and a photoperiod of 12 hours light and 12 hours dark.

### Microparticle bombardment

*Oryza sativa spp. japonica* cultivar Kitaake was used for transient transformation of leaf and root cells by microparticle bombardment. Seeds were de-husked prior to sterilisation in 5% (v/v) sodium hypochlorite solution containing 0.5% (v/v) Tween-20 for 30 minutes. Seeds were placed on half strength Murashige and Skoog (MS) medium containing 0.8% (w/v) agar and 1% (w/v) sucrose, and germinated in the dark at 28°C. Seedlings were grown in the dark at 28°C for 12 to 16 days and leaf 2 (of 12-day-old seedlings), leaf 3 (of 16-day-old seedlings), or roots used for microparticle bombardment.

Plasmid DNA was bound to 1 μm DNAdel™ Gold Carrier Particles (Seashell Technology) according to the manufacturer’s instructions. For each expression vector, 5 μg DNA were bound to 1.5 mg particles prior to resuspension in 70 μl 100% (v/v) ethanol. Seven μl of the particle suspension was transferred to plastic macrocarriers (Bio-Rad) and allowed to dry completely at room temperature. For each transformation, leaf blades or roots were cut into 4-cm-long pieces, placed on 0.8% (w/v) water agar and bombarded three times with a Bio-Rad PDS-1000/He particle delivery system using 1550 psi rupture disks (Bio-Rad). For transformation of leaf cells, the abaxial side of the leaf blade was bombarded. After bombardment, leaf blades or roots were flooded with half strength MS solution containing 1% (w/v) sucrose and incubated in the dark (or continuous light for chloroplast-targeted proteins) at 28°C for 24 hours before analysis by confocal scanning laser microscopy or staining for GUS.

### GUS staining

To visualise GUS activity in transiently transformed cells, leaf blades or roots were submerged in GUS staining solution [0.1 M Na_2_HPO_4_ pH 7.0, 0.5 mM K ferricyanide, 0.5 mM K ferrocyanide, 0.06% (v/v) Triton X-100, 10 mM Na_2_EDTA pH 8.0, 1 mM X-Gluc (5-bromo-4-chloro-3-indolyl-beta-D-glucuronide)] 24 hours after microparticle bombardment. For leaf blades, a vacuum was applied 4 times for 2 minutes each time to submerge the tissue. Plant material was incubated in GUS staining solution for 6 to 24 hours at 37°C in the dark and fixed in a 3:1 solution of ethanol and acetic acid for 30 minutes at room temperature. For leaf blades, chlorophyll was cleared with 70% (v/v) ethanol, followed by incubation in 5% (w/v) NaOH for 2 hours at 37°C. Brightfield images of GUS-stained plant tissue were obtained with an Olympus BX41 light microscope.

### Tissue preparation and confocal imaging

Prior to confocal imaging, leaf tissue was prepared by either scraping the leaf surface or clearing the leaves using the ClearSee protocol to enable visualisation of deeper tissue layers. The adaxial side of 4-cm-long segments of the leaf blade from mature rice leaves was scraped 2 to 3 times with a sharp razor blade, and transferred to Phosphate Buffered Saline (PBS) to avoid drying, and then mounted on a microscope slide with the scraped surface facing upwards. The ClearSee protocol was performed as previously described (Kurihara et al., 2015). Leaf blades were cut into 5-mm-long segments with a razor blade and fixed in 20 ml 4% (w/v) paraformaldehyde solution containing a small drop of silwet-L77 for 1 hour under vacuum at room temperature. During tissue fixation, vacuum was released and reapplied 5 times. After washing the leaves twice with PBS, they were cleared with ClearSee solution [10% (w/v) xylitol, 15% (w/v) sodium deoxycholate, 25% (w/v) urea] for 3 to 4 weeks with gentle agitation at room temperature. The ClearSee solution was replaced every 2 to 3 days for four weeks.

Confocal imaging was performed on a Leica TCS SP8 X using a 10X air objective (HC PL APO CS2 10X 0.4 Dry) with or without optical zoom, a hybrid detector for fluorescent protein detection and a PMT for detection of chlorophyll autofluorescence. The following excitation (Ex) and emission (Em) wavelengths were used for detection: mTurquoise2 (Ex = 442, Em = 471-481), mTFP1 (Ex = 442, Em = 477-517), mClover3 (Ex = 506, Em = 513-523), mNeonGreen (Ex = 506, Em = 512-522), mCitrine (Ex = 516, Em = 524-534), mYPet (Ex = 517, Em = 525-535), mKOκ (Ex = 551, Em = 558-568), tdTomato (Ex = 554, Em = 576-586), TagRFP-T (Ex = 555, Em = 579-589), mRuby3 (Ex = 558, Em = 587-597), mCardinal (Ex = 604, Em = 654-664), mKate2 (Ex = 588 nm, Em = 628-638), chlorophyll autofluorescence (Ex = 488, Em = 672-692).

## Results

### Identification of fluorescent reporters that are stably expressed and detectable in the rice leaf blade

Fluorescence imaging of rice leaves is impaired by high levels of light scattering arising from the leaf surface, which impedes in-focus, high-resolution imaging of deeper leaf layers such as the mesophyll cells (Qin et al., 2011; Clark and Lister, 1975); Fig. S2a,d). In recent years, several methods have been developed to clear tissue for whole-organ or whole-plant imaging (Palmer et al., 2015; Kurihara et al., 2015; Timmers, 2016). Amongst these, the ClearSee protocol enables deep imaging in Arabidopsis whole leaves while at the same time maintaining the stability of fluorescent proteins (Kurihara et al., 2015). We tested whether ClearSee treatment can also be used to improve whole-leaf imaging in rice. Four weeks of clearing rice leaves led to an improved visualisation of different layers of the leaf blade, including the mesophyll cells, similar to what has previously been shown in Arabidopsis (Fig. S2b,e). While these results suggest that ClearSee is an effective method for deep tissue imaging in rice leaves, the protocol requires fixation followed by several weeks of clearing. We therefore sought a faster method that allows live imaging of the leaf interior. We manually removed the leaf surface by scraping the adaxial side of each leaf with a razor blade and mounted the leaf with the scraped surface facing upwards to allow unimpeded laser interrogation of the leaf interior. We found that this enabled in-focus visualisation of the mesophyll cells (Fig. S2c,f). However, we noticed the presence of bright autofluorescent structures in the mesophyll layer in almost all regions of the visible spectrum (Fig. S2i,l,o), which were also present in non-scraped leaves (Fig. S2g,j,m), but appeared to be less prominent in ClearSee-treated leaves (Fig. S2h,k,n). Together, these results suggest that both methods can be used to overcome light scattering for high resolution imaging of the leaf interior. While scraping the leaf surface is a quick method to improve imaging, the ClearSee protocol requires several weeks of clearing, but has the advantage of reducing signal from autofluorescent structures in deeper tissue layers.

Over the years, a large number of fluorescent protein variants with different excitation and emission spectra have been developed and successfully used in many plant species. Being able to combine these different fluorescent reporters allows the co-visualisation of multiple compartments or biological processes in the same cell (Shaner et al., 2005; DeBlasio et al., 2010). However, to our knowledge only GFP has been visualised in the rice leaf blade (Johnson et al., 2005; Reynoso et al., 2018). To expand the range of fluorescent proteins that can be used for functional analysis in rice leaves, we assessed twelve proteins that emit light from the cyan to far-red regions of the spectrum for inherent brightness in stably transformed rice plants (Figure S1). To facilitate imaging and clearly distinguish genuine fluorescent protein signal from autofluorescent structures, we placed each reporter under the control of the maize *PEPC* promoter to direct gene expression to mesophyll cells (Matsuoka et al., 1994) and used a nuclear localisation signal to specifically label nuclei. We assessed five to nine independent T_0_ lines for each reporter and scraped the leaf surface prior to confocal microscopy for improved visualisation. The degree of signal from each protein was variable, but five were robustly detectable despite autofluorescent structures in the mesophyll layer (Figure 1, Figure S3, Table 1). mTurquoise2 generated the clearest signal, but mClover3, mNeonGreen, mKOκ, and tdTomato were also clearly detectable (Figure 1). To confirm that expression of these fluorescent proteins is stable in subsequent generations of transgenic rice plants, three independent T_1_ plants for each construct were imaged by confocal microscopy. T_1_ plants showed detectable signal in mesophyll nuclei for all five proteins (Figure S4), indicating that transgene expression and fluorescent protein accumulation were maintained in the next generation of transgenic plants.

**Figure 1:**
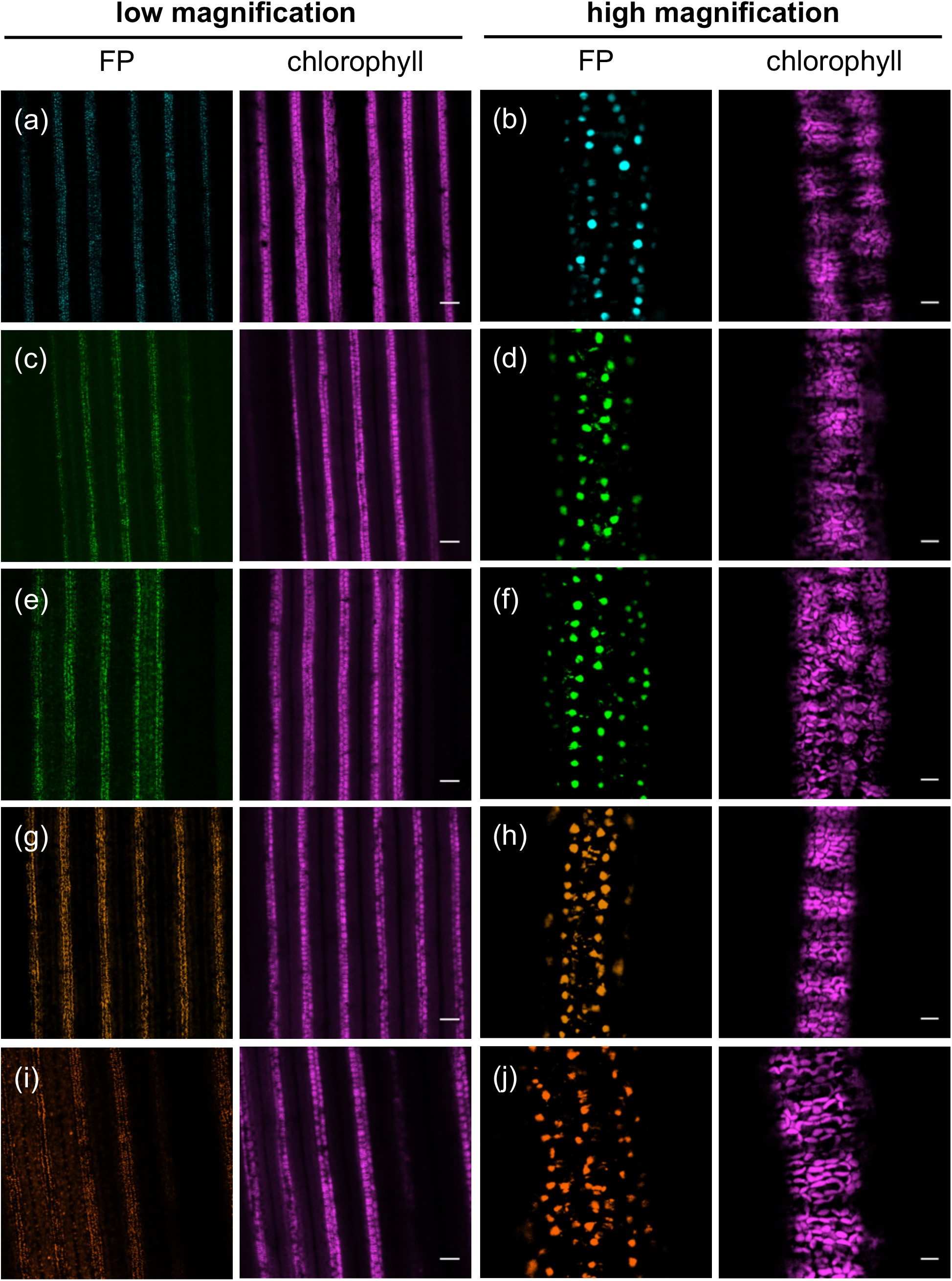
Fluorescent proteins expressed in mesophyll cells of stably transformed T_0_ rice plants. (a) and (b) T_0_ plants expressing *ZmPEPC*_*pro*_:*mTurquoise2-NLS*. (c) and (d) T_0_ plants expressing *ZmPEPC*_*pro*_:*mClover3-NLS*. (e) and (f) T_0_ plants expressing *ZmPEPC*_*pro*_:*mNeonGreen-NLS*. (g) and (h) T_0_ plants expressing *ZmPEPC*_*pro*_:*mKOκ-NLS*. (i) and (j) T_0_ plants expressing *ZmPEPC*_*pro*_:*tdTomato-NLS*. (a), (c), (e), (g), and (i) show each fluorescent protein (FP). (b), (d), (f), (h), and (j) show chlorophyll. High magnification images, scale bars represent 10 µm. Low magnification images, scale bars represent 100 µm.

We also tested whether the stability of the five fluorescent reporters is maintained after ClearSee treatment. To this end, leaf tissue from transgenic plants was fixed with paraformaldehyde. After four weeks of ClearSee treatment, a strong nuclear fluorescent signal was detectable in leaves expressing mTurquoise2, mKOκ, and tdTomato (Figure S5a-c, j-o). However, no nuclear signal was detectable in mesophyll cells of rice plants expressing mClover3 and mNeonGreen, suggesting that ClearSee treatment negatively affects the stability of mClover3 and mNeonGreen in rice leaves under the conditions tested here (Figure S5d-i).

### A transient expression system for functional analysis in the rice leaf blade

As an alternative to the time-consuming and labour-intensive process required for stable transformation of rice, heterologous expression systems or protoplast assays are routinely used to investigate the regulation of rice genes as well as the function and subcellular localisation of rice proteins. Although these techniques can be used to assess gene and protein function in a relatively short amount of time, they do not allow analysis in cells of intact rice tissues. Microparticle bombardment has successfully been employed to transform individual cells in whole leaves of several plant species (Brown et al., 2011; Reyna-Llorens et al., 2018). For rice, even though particle bombardment has been used for transient expression in epidermal cells of the leaf sheath (Dangol et al., 2017; Meng et al., 2014; Wang et al., 2018), there are very few reports of successful microparticle bombardment of the mature rice leaf blade (Jia et al., 2000). We tested whether microparticle bombardment can be used as a fast and simple technique to transiently transform different cell types in the leaf blade of fully developed rice leaves. Epidermal, guard, trichome, bulliform, mesophyll, bundle sheath, mestome sheath, and veinal cells were successfully transformed after bombardment with a *GUS* reporter gene under the control of the constitutively active *CaMV35S* promoter (Figure 2a-h). On average, 34 ± 18 cells per leaf (n = 6 leaves) showed transgene expression, suggesting that microparticle bombardment is an efficient method to transiently transform a range of different cell types, and importantly cells within the leaf interior, in intact, fully expanded rice leaves. We next tested whether microparticle bombardment is suitable to fluorescently label different cell compartments simultaneously using the fluorescent reporters that we identified in our screen. To this end, vectors for the co-expression of mTurquoise2 and mNeonGreen targeted to either the nucleus (Cutler et al., 2000), plasma membrane (Dangol et al., 2017), or chloroplast (Shen et al., 2017) were generated. Bombarding rice leaves with these vectors resulted in transformed cells containing both mTurquoise2 and mNeonGreen targeted to two different cell compartments (Figure 3). Similar to the results with the *GUS* reporter gene, we were able to detect fluorescent proteins using confocal microscopy in a range of different cell types, including epidermal cells and mesophyll cells (Figure 3). In addition to using microparticle bombardment for rice leaves, we also tested whether tissues such as roots could be transformed using our experimental conditions. We found that bombardment of roots with a *GUS* reporter gene resulted in successfully transformed root cells (Figure 2i). Moreover, the nuclei and plasma membrane of root cells could be simultaneously labelled with mTurquoise2 and mNeonGreen (Figure S6). These results indicate that mTurquoise2 and mNeonGreen are not only suitable for imaging in rice leaves, but are also stably expressed and visible in rice roots.

**Figure 2:**
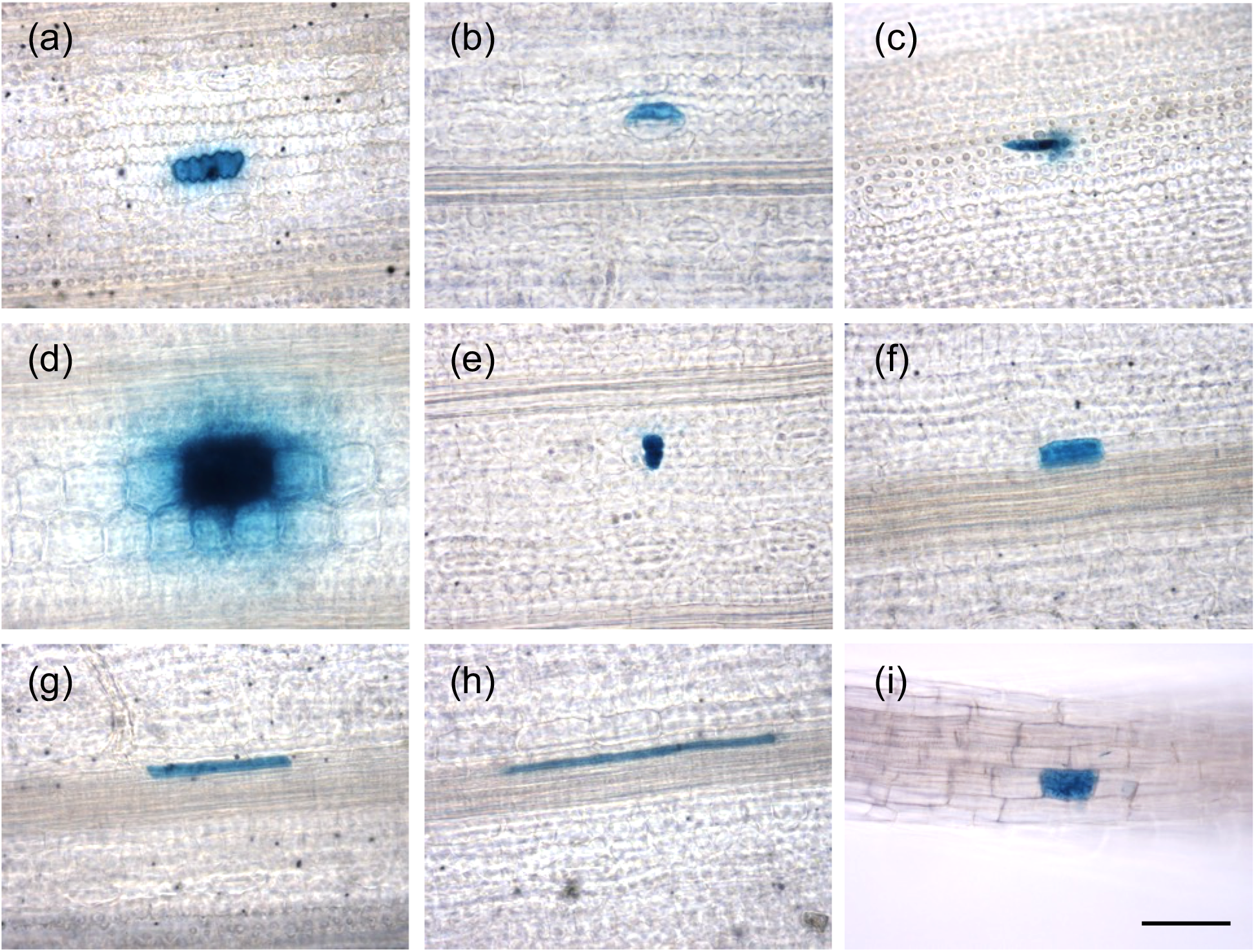
Microparticle bombardment of rice leave blades and roots. GUS accumulation in leaf and root cells transformed with a *CaMV35S*_*pro*_:*GUS* vector using microparticle bombardment. Transformed epidermal (a), guard (b), trichome (c), bulliform (d), mesophyll (e), bundle sheath (f), mestome sheath (g), veinal (h), and root cell (i). Scale bars represent 50 µm.

**Figure 3:**
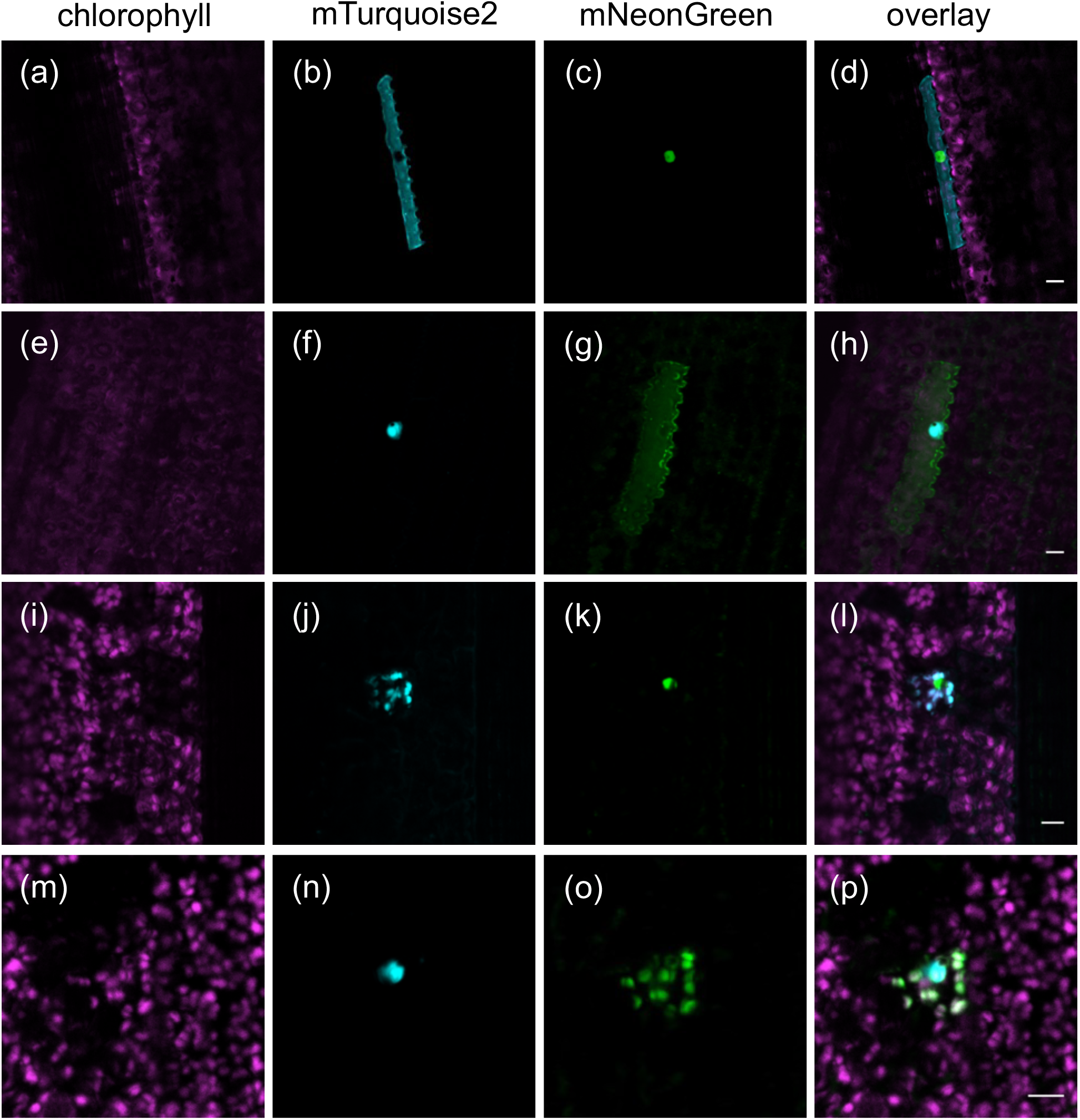
Fluorescent proteins targeted to different cell compartments in transiently transformed rice leaf cells. (a)-(d) Epidermal cell transformed with a construct expressing plasma membrane-localised mTurquoise2 and nuclear-localised mNeonGreen. (e)-(h) Epidermal cell transformed with a construct expressing nuclear-localised mTurquoise2 and plasma membrane-localised mNeonGreen. (i)-(l) Mesophyll cell transformed with a construct expressing chloroplast-localised mTurquoise2 and nuclear-localised mNeonGreen. (m)-(p) Mesophyll cell transformed with a construct expressing nuclear-localised mTurquoise2 and chloroplast-localised mNeonGreen. Scale bars represent 10 µm.

## Discussion

Rice is one of the most important food crops in the world. If we aim to engineer rice with increased yield, improvements in our understanding of its biology are likely required. Fluorescent reporters significantly facilitate the analysis of biological processes; however, such tools are currently limited for rice leaves. Of the twelve fluorescent proteins that we tested, seven could not be detected reliably under our experimental conditions. Several reasons could account for this observation. It is possible that codon optimization introduces sequence motifs that repress either transcription or translation, or that it adversely affects mRNA stability or interferes with protein folding and thus disrupts protein function (Hanson and Coller, 2017; Yu et al., 2015; Zhao et al., 2017). It is also possible that transcripts are silenced in transgenic plants, or that proteins are unstable and degraded. Further analyses will be required to test these possibilities. In addition to EGFP, which has been detected in epidermal and guard cells of the rice leaf blade (Reynoso et al., 2018), we have identified an additional five fluorescent reporters that emit light in the cyan, green, orange and red regions of the spectrum and are robustly detectable in the rice leaf blade. Of these reporters, mTurquoise2 generated the most clearly detectable signal owing to its brightness and the low prevalence of autofluorescent structures emitting in the cyan region of the spectrum. Although the prevalence of autofluorescent structures in the green to orange regions of the spectrum appears to be higher, the fluorescent proteins mClover3, mNeonGreen, mKOκ, and tdTomato were also visible. Based on these results, we propose that these five fluorescent reporters are suitable for fluorescence imaging and other fluorescence-based applications, such as protein tagging for immunoprecipitation or fluorescence-activated cell sorting, in rice leaves. Because the identified proteins emit light in three different spectral ranges (cyan, green and orange/red), combinations of these reporters can be used to co-visualise several proteins and/or cell compartments in leaves and other organs of rice.

We have shown that ClearSee treatment is compatible with use of mTurquoise2, mKOκ, and tdTomato in rice leaves. However, no nuclear-localised fluorescence was detected in leaves expressing mClover3 or mNeonGreen after treating tissues with ClearSee. The fluorescent protein mClover3 has previously successfully been detected in cleared, whole Arabidopsis leaves (Kurihara et al., 2015). Our inability to detect mClover3 and mNeonGreen in cleared rice leaves could be due to inherent differences between rice and Arabidopsis; however, it seems more likely that the detergent we used during fixation of rice, which was not used during fixation of Arabidopsis, adversely affects protein stability or fluorescence of mClover3 and mNeonGreen.

Because stable transformation of rice is a time-consuming process, heterologous expression systems or protoplast assays are used to test the function of rice proteins in a time-efficient manner. Microparticle bombardment for transient transformation of cells in intact rice tissues such as leaves or roots is a useful alternative to these techniques as it has several advantages. The preparation of microparticles and the bombardment of plant tissue can be performed in a short amount of time and protein expression is detectable in a relatively high number of cells only 24 hours after bombardment, making this a fast and efficient transformation method. In addition, it ensures that protein function is analysed in the relevant cell and tissue context. Microparticle bombardment is not only suitable for analysing processes such as the subcellular localization of proteins in different cell types of rice, but can also be used to study protein-protein interactions or the spatial regulation of gene expression. Together with a high-throughput cloning technique such as Golden Gate (Engler et al., 2009; Patron et al., 2015), the findings reported here should facilitate molecular and cell biology analysis in rice, for example by allowing the rapid testing of gene expression vectors prior to stable transformation, analysis of protein-transcription factor binding through transactivation assays, protein-protein interactions by bimolecular fluorescence complementation, and subcellular targeting of proteins.

## Acknowledgements

The work was supported by a C_4_ Rice project grant from The Bill and Melinda Gates Foundation to the University of Oxford (2015-2019) and sLola BB/P003117/1 from the BBSRC.

## Authors contributions

J.M.H, L.H.L., and S.E. designed the experimental work. L.H.L. and S.E. cloned expression vectors for transient and stable transformation. N.W. carried out stable transformation of rice. L.H.L. performed microparticle bombardment experiments. L.H.L., S.E., and N.W. performed the microscopy. L.H.L., S.E and J.M.H. wrote the manuscript. All authors revised the manuscript and approved the final version.

## References

Abiko, M., Furuta, K., Yamauchi, Y., Fujita, C., Taoka, M., Isobe, T., and Okamoto, T. (2013). Identification of Proteins Enriched in Rice Egg or Sperm Cells by Single-Cell Proteomics. PLoS One 8.

Ai, H., Henderson, J.N., Remington, S.J., and Campbell, R.E. (2006). Directed evolution of a monomeric, bright and photostable version of Clavularia cyan fluorescent protein: structural characterization and applications in fluorescence imaging. Biochem. J. 400: 531–40.

Bajar, B.T., Wang, E.S., Lam, A.J., Kim, B.B., Jacobs, C.L., Howe, E.S., Davidson, M.W., Lin, M.Z., and Chu, J. (2016). Improving brightness and photostability of green and red fluorescent proteins for live cell imaging and FRET reporting. Sci. Rep. 6: 20889.

Bellucci, M., De Marchis, F., Mannucci, R., Bock, R., and Arcioni, S. (2005). Cytoplasm and chloroplasts are not suitable subcellular locations for β-zein accumulation in transgenic plants. J. Exp. Bot. 56: 1205–1212.

Bennetzen, J.L. and Freeling, M. (1997). The unified grass genome: Synergy in synteny. Genome Res. 7: 301–306.

Birnbaum, K., Jung, J.W., Wang, J.Y., Lambert, G.M., Hirst, J.A., Galbraith, D.W., and Benfey, P.N. (2005). Cell type-specific expression profiling in plants via cell sorting of protoplasts from fluorescent reporter lines. Nat. Methods 2: 615–619.

Brown, N.J., Newell, C.A., Stanley, S., Chen, J.E., Perrin, A.J., Kajala, K., and Hibberd, J.M. (2011). Independent and Parallel Recruitment of Preexisting Mechanisms Underlying C4 Photosynthesis. Science 331: 1436–1439.

Chu, J. et al. (2014). Non-invasive intravital imaging of cellular differentiation with a bright red-excitable fluorescent protein. Nat. Methods 11: 572.

Chu, T.T.H. et al. (2018). Sub-cellular markers highlight intracellular dynamics of membrane proteins in response to abiotic treatments in rice. Rice 11.

Clark, J.B. and Lister, G.R. (1975). Photosynthetic Action Spectra of Trees. Plant Physiol. 55: 407–413.

Cornejo, M.J., Luth, D., Blankenship, K.M., Anderson, O.D., and Blechl, A.E. (1993). Activity of a maize ubiquitin promoter in transgenic rice. Plant Mol. Biol. 23: 567–581.

Cutler, S.R., Ehrhardt, D.W., Griffitts, J.S., and Somerville, C.R. (2000). Random GFP∷cDNA fusions enable visualization of subcellular structures in cells of Arabidopsis at a high frequency. Proc. Natl. Acad. Sci. 97: 3718 LP – 3723.

Dangol, S., Singh, R., Chen, Y., and Jwa, N. (2017). Molecules and Cells Visualization of Multicolored in vivo Organelle Markers for Co-Localization Studies in Oryza sativa. Mol. Cells 40: 828–836.

DeBlasio, S.L., Sylvester, A.W., and Jackson, D. (2010). Illuminating plant biology: Using fluorescent proteins for high-throughput analysis of protein localization and function in plants. Brief. Funct. Genomics 9: 129–138.

Engler, C., Gruetzner, R., Kandzia, R., and Marillonnet, S. (2009). Golden gate shuffling: a one-pot DNA shuffling method based on type IIs restriction enzymes. PLoS One 4: e5553.

De Folter, S., Urbanus, S.L., Van Zuijlen, L.G.C., Kaufmann, K., and Angenent, G.C. (2007). Tagging of MADS domain proteins for chromatin immunoprecipitation. BMC Plant Biol. 7: 1–11.

Goedhart, J., Von Stetten, D., Noirclerc-Savoye, M., Lelimousin, M., Joosen, L., Hink, M.A., Van Weeren, L., Gadella, T.W.J., and Royant, A. (2012). Structure-guided evolution of cyan fluorescent proteins towards a quantum yield of 93%. Nat. Commun. 3.

Goff, S.A. et al. (2002). A Draft Sequence of the Rice Genome (Oryza sativa L. ssp. japonica). Science 296: 92 LP – 100.

Griesbeck, O., Baird, G.S., Campbell, R.E., Zacharias, D.A., and Tsien, R.Y. (2001). Reducing the Environmental Sensitivity of Yellow Fluorescent Protein: MECHANISM AND APPLICATIONS. J. Biol. Chem. 276: 29188–29194.

Gupta, P., Rangan, L., Ramesh, T.V., and Gupta, M. (2016). Comparative analysis of contextual bias around the translation initiation sites in plant genomes. J. Theor. Biol. 404: 303–311.

Hanson, G. and Coller, J. (2017). Codon optimality, bias and usage in translation and mRNA decay. Nat. Rev. Mol. Cell Biol. 19: 20.

Hanson, M.R. and Köhler, R.H. (2001). GFP imaging: Methodology and application to investigate cellular compartmentation in plants. J. Exp. Bot. 52: 529–539.

Haseloff, J. (1998). GFP Variants for Multispectral Imaging of Living Cells. In Methods in Cell Biology (Academic Press), pp. 139–151.

Haseloff, J., Siemering, K.R., Prasher, D.C., and Hodge, S. (1997). Removal of a cryptic intron and subcellular localization of green fluorescent protein are required to mark transgenic Arabidopsis plants brightly. Proc. Natl. Acad. Sci. U. S. A. 94: 2122–2127.

He, C., Lin, Z., McElroy, D., and Wu, R. (2009). Identification of a rice Actin2 gene regulatory region for high-level expression of transgenes in monocots. Plant Biotechnol. J. 7: 227–239.

Hebsgaard, S.M., Korning, P.G., Tolstrup, N., Engelbrecht, J., Rouzé, P., and Brunak, S. (1996). Splice site prediction in Arabidopsis thaliana pre-mRNA by combining local and global sequence information. Nucleic Acids Res. 24: 3439–52.

Hiei, Y. and Komari, T. (2008). Agrobacterium-mediated transformation of rice using immature embryos or calli induced from mature seed. Nat. Protoc. 3: 824–834.

Jefferson, R.A., Kavanagh, T.A., and Bevan, M.W. (1987). GUS fusions: beta-glucuronidase as a sensitive and versatile gene fusion marker in higher plants. EMBO J. 6: 3901–3907.

Jia, Y., Mcadams, S.A., Bryan, G.T., Hershey, H.P., and Valent, B. (2000). Direct interaction of resistance gene and avirulence gene products confers rice blast resistance. 19: 4004–4014.

Johnson, A.A.T., Hibberd, J.M., Gay, C., Essah, P.A., Haseloff, J., Tester, M., and Guiderdoni, E. (2005). Spatial control of transgene expression in rice (Oryza sativa L.) using the GAL4 enhancer trapping system. Plant J. 41: 779–789.

Joshi, C.P., Zhou, H., Huang, X., and Chiang, V.L. (1997). Context sequences of translation initiation codon in plants. Plant Mol. Biol. 35: 993–1001.

Kim, J.Y., Yuan, Z., Cilia, M., Khalfan-Jagani, Z., and Jackson, D. (2002). Intercellular trafficking of a *KNOTTED1* green fluorescent protein fusion in the leaf and shoot meristem of *Arabidopsis*. Proc. Natl. Acad. Sci. 99: 4103–4108.

Kitajima, A., Asatsuma, S., Okada, H., Hamada, Y., Kaneko, K., Nanjo, Y., Kawagoe, Y., Toyooka, K., Matsuoka, K., Takeuchi, M., Nakano, A., and Mitsui, T. (2009). The Rice alpha-Amylase Glycoprotein Is Targeted from the Golgi Apparatus through the Secretory Pathway to the Plastids. Plant Cell 21: 2844–2858.

Kobae, Y. and Hata, S. (2010). Dynamics of periarbuscular membranes visualized with a fluorescent phosphate transporter in arbuscular mycorrhizal roots of rice. Plant Cell Physiol. 51: 341–353.

Kurihara, D., Mizuta, Y., Sato, Y., and Higashiyama, T. (2015). ClearSee: a rapid optical clearing reagent for whole-plant fluorescence imaging. Development 142: 4168–4179.

Lam, S.K., Siu, C.L., Hillmer, S., Jang, S., An, G., Robinson, D.G., and Jiang, L. (2007). Rice SCAMP1 Defines Clathrin-Coated, trans-Golgi-Located Tubular-Vesicular Structures as an Early Endosome in Tobacco BY-2 Cells. Plant Cell 19: 296–319.

Luehrsen, K.R., de Wet, J.R., and Walbot, V. (1992). Transient expression analysis in plants using firefly luciferase reporter gene. Methods Enzymol. 216: 397–414.

Matsumoto, T. et al. (2005). The map-based sequence of the rice genome. Nature 436: 793–800.

Matsuoka, M., Kyozuka, J., Shimamoto, K., and Kano-Murakami, Y. (1994). The promoters of two carboxylases in a C4 plant (maize) direct cell-specific, light-regulated expression in a C3 plant (rice). Plant J. 6: 311–319.

Meng, W., Hsiao, A.S., Gao, C., Jiang, L., and Chye, M.L. (2014). Subcellular localization of rice acyl-CoA-binding proteins (ACBPs) indicates that OsACBP6::GFP is targeted to the peroxisomes. New Phytol. 203: 469–482.

Mohanty, A., Luo, A., DeBlasio, S., Ling, X., Yang, Y., Tuthill, D.E., Williams, K.E., Hill, D., Zadrozny, T., Chan, A., Sylvester, A.W., and Jackson, D. (2009). Advancing Cell Biology and Functional Genomics in Maize Using Fluorescent Protein-Tagged Lines. Plant Physiol. 149: 601–605.

Nakagawa, S., Niimura, Y., Gojobori, T., Tanaka, H., and Miura, K. ichiro (2008). Diversity of preferred nucleotide sequences around the translation initiation codon in eukaryote genomes. Nucleic Acids Res. 36: 861–871.

Nguyen, A.W. and Daugherty, P.S. (2005). Evolutionary optimization of fluorescent proteins for intracellular FRET. Nat. Biotechnol. 23: 355–360.

Palmer, W.M., Martin, A.P., Flynn, J.R., Reed, S.L., White, R.G., Furbank, R.T., and Grof, C.P.L. (2015). PEA-CLARITY: 3D molecular imaging of whole plant organs. Sci. Rep. 5: 1–6.

Patron, N.J. et al. (2015). Standards for plant synthetic biology: A common syntax for exchange of DNA parts. New Phytol. 208: 13–19.

Prasher, D.C., Eckenrode, V.K., Ward, W.W., Prendergast, F.G., and Cormier, M.J. (1992). Primary structure of the Aequorea victoria green-fluorescent protein. Gene 111: 229–233.

Qin, B.X., Tang, D., Huang, J., Li, M., Wu, X.R., Lu, L.L., Wang, K.J., Yu, H.X., Chen, J.M., Gu, M.H., and Cheng, Z.K. (2011). Rice OsGL1-1 is involved in leaf cuticular wax and cuticle membrane. Mol. Plant 4: 985–995.

Reyna-Llorens, I., Burgess, S.J., Reeves, G., Singh, P., Stevenson, S.R., Williams, B.P., Stanley, S., and Hibberd, J.M. (2018). Ancient duons may underpin spatial patterning of gene expression in C 4 leaves. Proc. Natl. Acad. Sci. 115: 1931–1936.

Reynoso, M.A., Pauluzzi, G.C., Kajala, K., Cabanlit, S., Velasco, J., Bazin, J., Deal, R., Sinha, N.R., Brady, S.M., and Bailey-Serres, J. (2018). Nuclear Transcriptomes at High Resolution Using Retooled INTACT. Plant Physiol. 176: 270 LP – 281.

Shaner, N.C., Campbell, R.E., Steinbach, P.A., Giepmans, B.N.G., Palmer, A.E., and Tsien, R.Y. (2004). Improved monomeric red, orange and yellow fluorescent proteins derived from Discosoma sp. red fluorescent protein. Nat. Biotechnol. 22: 1567–1572.

Shaner, N.C., Lambert, G.G., Chammas, A., Ni, Y., Paula, J., Baird, M.A., Sell, B.R., Allen, J.R., Day, R.N., Davidson, M.W., and Wang, J. (2013). A bright monomeric green fluorescent protein derived from Branchiostoma lanceolatum Nathan. Nat. Methods 10: 407–409.

Shaner, N.C., Lin, M.Z., McKeown, M.R., Steinbach, P.A., Hazelwood, K.L., Davidson, M.W., and Tsien, R.Y. (2008). Improving the photostability of bright monomeric orange and red fluorescent proteins. Nat. Methods 5: 545.

Shaner, N.C., Steinbach, P.A., and Tsien, R.Y. (2005). A guide to choosing fluorescent proteins. Nat. Methods 2: 905–909.

Shcherbo, D. et al. (2009). Far-red fluorescent tags for protein imaging in living tissues. Biochem. J. 418: 567 LP – 574.

Shen, B.R., Zhu, C.H., Yao, Z., Cui, L.L., Zhang, J.J., Yang, C.W., He, Z.H., and Peng, X.X. (2017). An optimized transit peptide for effective targeting of diverse foreign proteins into chloroplasts in rice. Sci. Rep. 7.

Supartana, P., Shimizu, T., Shioiri, H., Nogawa, M., Nozue, M., and Kojima, M. (2005). Development of Simple and Efficient in Planta Transformation Method for Rice (Oryza sativa L.) Using Agrobacterium tumefaciens. J. Biosci. Bioeng. 100: 391–397.

Timmers, A.C.J. (2016). Light microscopy of whole plant organs. J. Microsc. 263: 165–170.

Toki, S., Hara, N., Ono, K., Onodera, H., Tagiri, A., Oka, S., and Tanaka, H. (2006). Early infection of scutellum tissue with Agrobacterium allows high-speed transformation of rice. Plant J. 47: 969–976.

Tsutsui, H., Karasawa, S., Okamura, Y., and Miyawaki, A. (2008). Improving membrane voltage measurements using FRET with new fluorescent proteins. Nat. Methods 5: 683.

Wang, H., Zhao, Q., Fu, J., Wang, X., and Jiang, L. (2018). Re-assessment of biolistic transient expression: An efficient and robust method for protein localization studies in seedling-lethal mutant and juvenile plants. Plant Sci. 274: 2–7.

Weber, E., Engler, C., Gruetzner, R., Werner, S., and Marillonnet, S. (2011). A modular cloning system for standardized assembly of multigene constructs. PLoS One 6: e16765.

Wu, T.M., Lin, K.C., Liau, W.S., Chao, Y.Y., Yang, L.H., Chen, S.Y., Lu, C.A., and Hong, C.Y. (2016). A set of GFP-based organelle marker lines combined with DsRed-based gateway vectors for subcellular localization study in rice (Oryza sativa L.). Plant Mol. Biol. 90: 107–115.

Yu, C.-H., Dang, Y., Zhou, Z., Wu, C., Zhao, F., Sachs, M.S., and Liu, Y. (2015). Codon Usage Influences the Local Rate of Translation Elongation to Regulate Co-translational Protein Folding. Mol. Cell 59: 744–754.

Zhang, Y., Su, J., Duan, S., Ao, Y., Dai, J., Liu, J., Wang, P., Li, Y., Liu, B., Feng, D., Wang, J., and Wang, H. (2011). A highly efficient rice green tissue protoplast system for transient gene expression and studying light/chloroplast-related processes. Plant Methods 7: 30.

Zhao, F., Yu, C.-H., and Liu, Y. (2017). Codon usage regulates protein structure and function by affecting translation elongation speed in Drosophila cells. Nucleic Acids Res. 45: 8484–8492.

