## Supplemental Figures for "Fluorescent reporters for functional analysis in rice leaves"

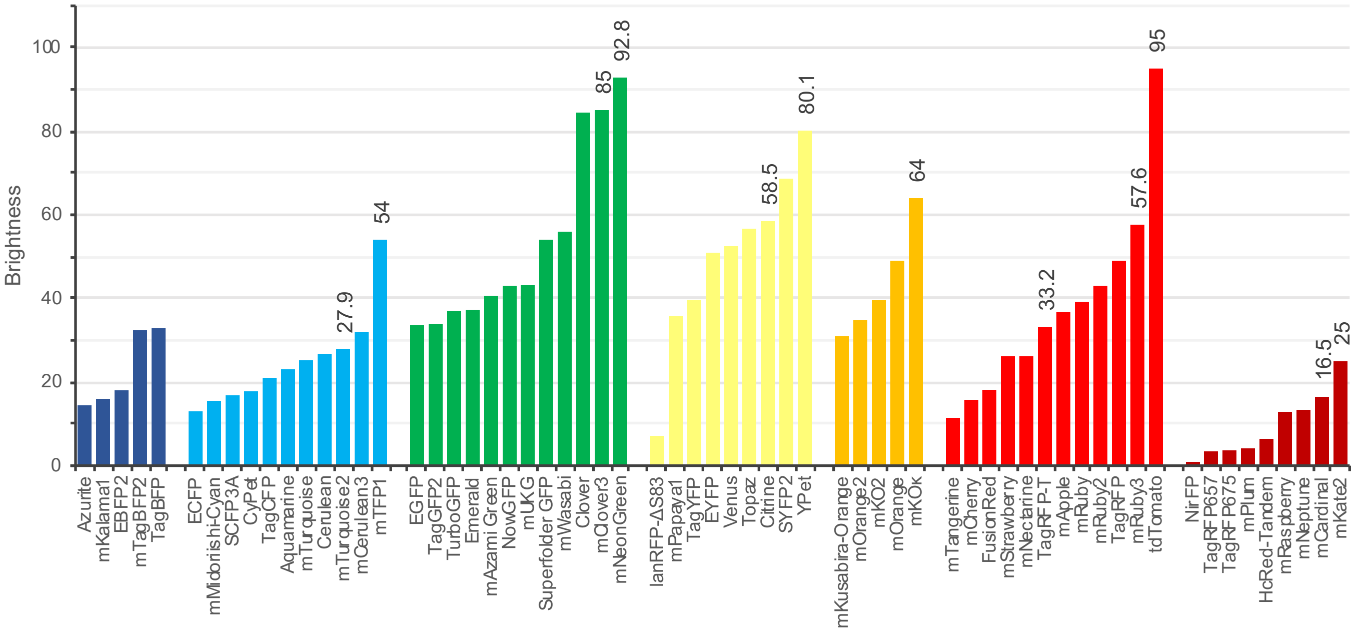


**Supplemental Figure 1: Fluorescent proteins chosen as candidate reporters to test in rice.** The twelve fluorescent proteins selected for screening are annotated above each bar with a number representing their relative brightness. From left to right: blue, cyan, green, yellow, orange, red and far red fluorescent proteins. With the exception of tdTomato, the displayed set of proteins was obtained from (http://www.fpvis.org/FP.html).

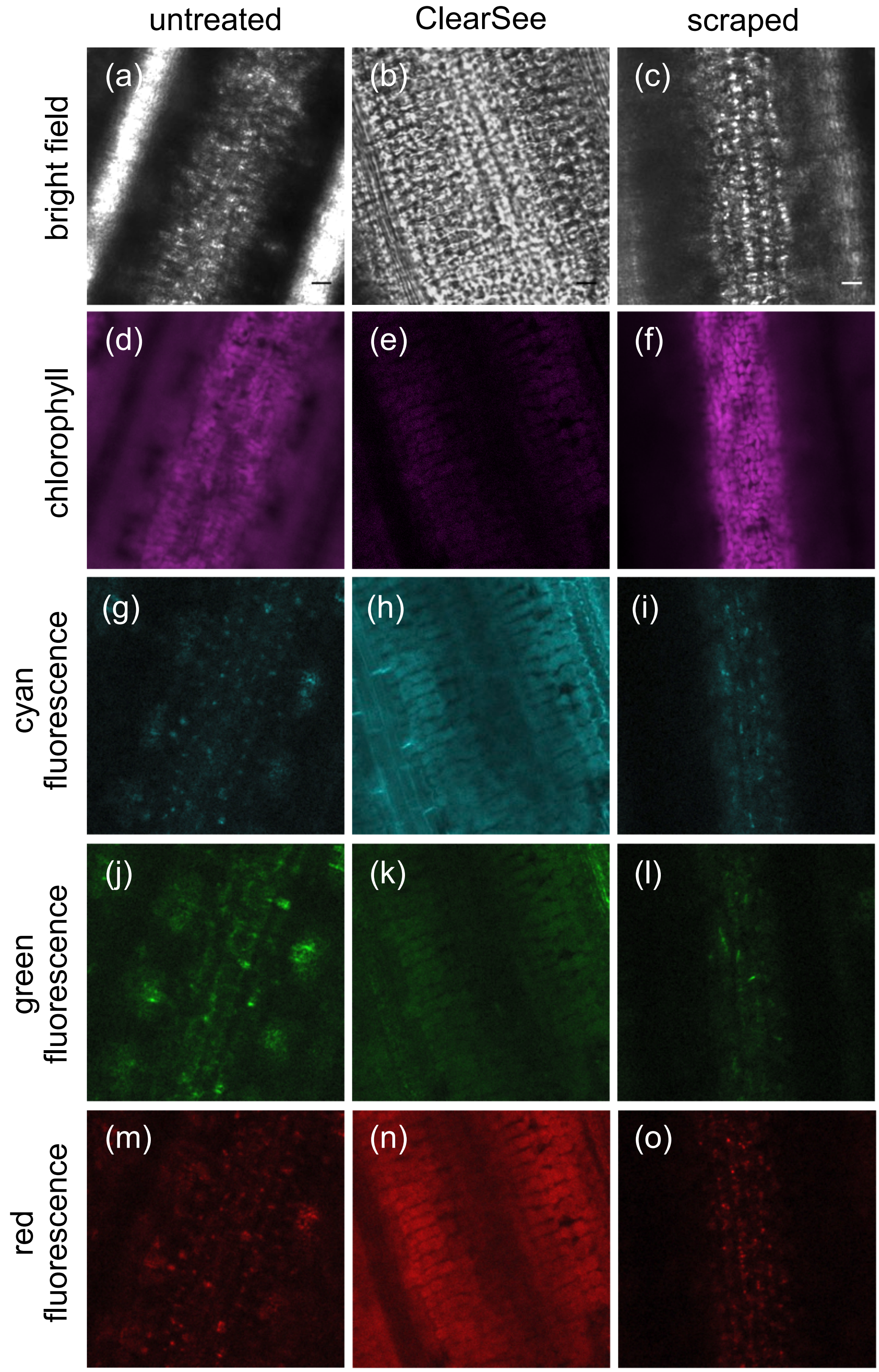


**Supplemental Figure 2: Deep imaging in the rice leaf blade.** Representative confocal images of the mesophyll layer from untreated wild-type leaves (a), (d), (g), (j) and (m). Note the lack of clarity due to light scattering in bright field (a) and chlorophyll (d) images. In contrast, bright-field imaging of mesophyll cells was facilitated with ClearSee (b), (e), (h), (k) and (n) or scraping (c), (f), (i), (l) and (o). Scraping also allowed distinct foci due to the chlorophyll autofluorescence from chloroplasts to be imaged (f). Autofluorescence in the cyan, green and red spectra were detected with mTurquoise2, mClover3 and mRuby3 settings, respectively. A gain of 250 was used to acquire images in the cyan, green and red spectra, except for (h), where a reduced gain of 100 was used due to high levels of background fluorescence in leaves treated with ClearSee. Scale bars represent 10 µm.

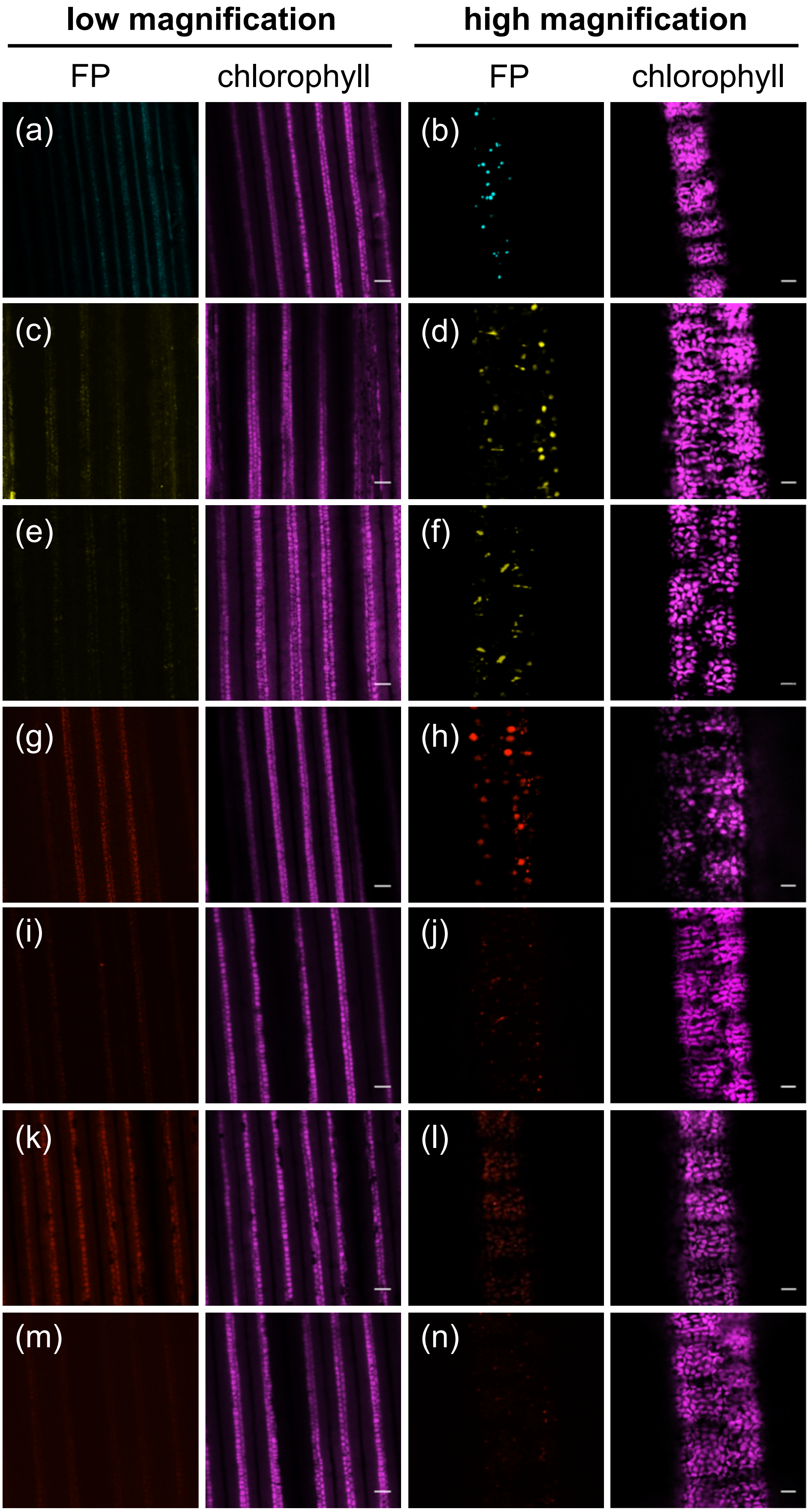


**Supplemental Figure 3: Fluorescent proteins expressed in mesophyll cells of stably transformed $T_0$ rice plants.** (a) and (b) $T_0$ plants expressing *ZmPEPC$_{pro}$:mTFP1-NLS.* (c) and (d) $T_0$ plants expressing *ZmPEPC$_{pro}$:mCitrine-NLS.* (e) and (f) $T_0$ plants expressing *ZmPEPC$_{pro}$:mYPet-NLS.* (g) and (h) $T_0$ plants expressing *ZmPEPC$_{pro}$:TagRFPT-NLS*. (i) and (j) $T_0$ plants expressing *ZmPEPC$_{pro}$:mRuby-NLS.* (k) and (l) $T_0$ plants expressing *ZmPEPC$_{pro}$:mCardinal-NLS.* (m) and (n) $T_0$ plants expressing *ZmPEPC$_{pro}$:mKate2-NLS*. (a), (c), (e), (g), (i), (k) and (m) show each fluorescent protein (FP). (b), (d), (f), (h), (j), (l) and (n) show chlorophyll. High magnification images, scale bars represent 10 μm. Low magnification images, scale bars represent 100 μm.

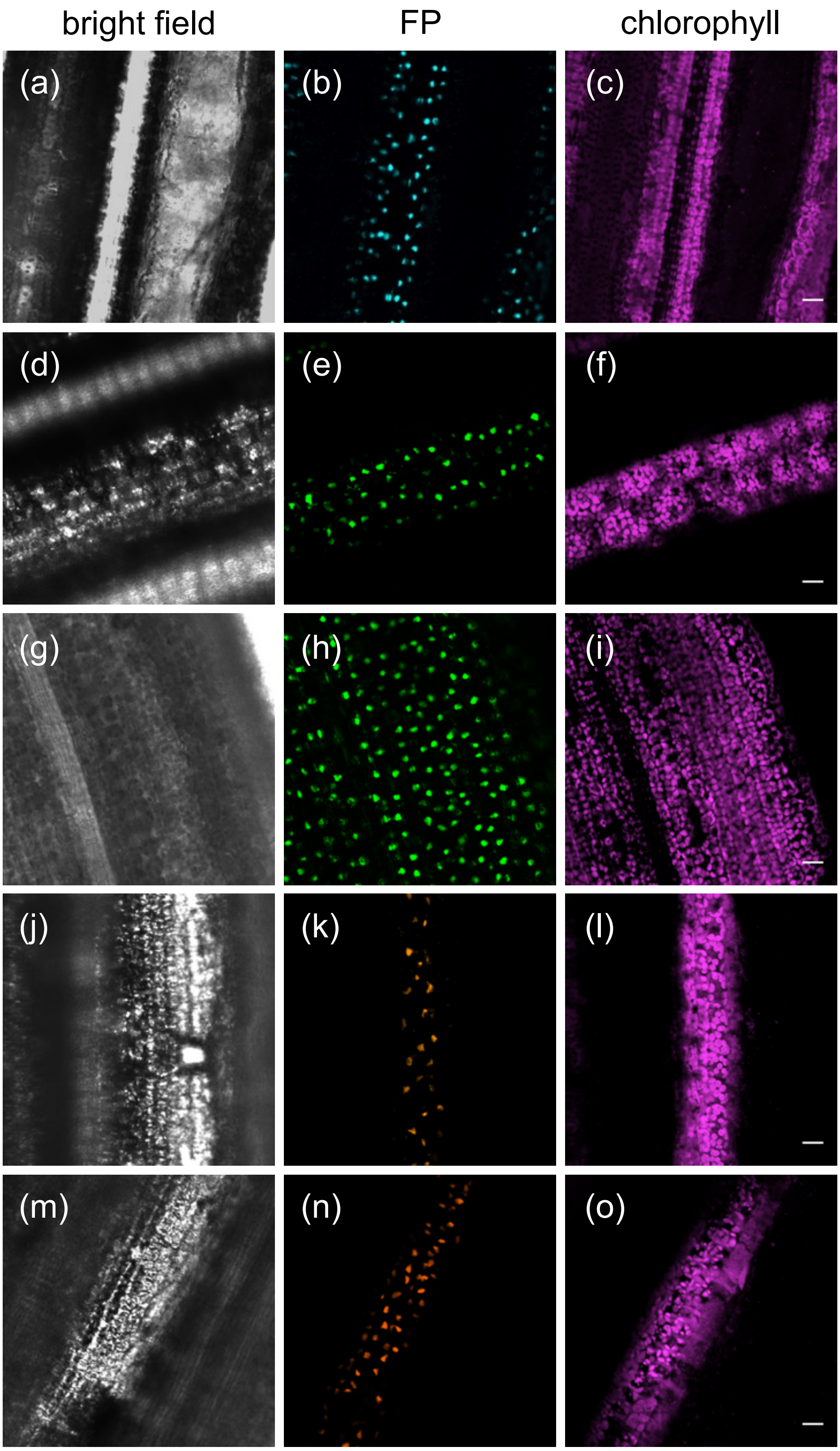


**Supplemental Figure 4: Fluorescent proteins expressed in mesophyll cells of stably transformed $T_1$ rice plants.** (a)-(c) $T_1$ plant expressing *ZmPEPCpro:mTurquoise2-NLS.* (d)-(f) $T_1$ plant expressing *$ZmPEPC_{pro}$:mClover3-NLS.* (g)-(i) $T_1$ plant expressing *$ZmPEPC_{pro}$:mNeonGreen-NLS.* (j)-(l) $T_1$ plant expressing *$ZmPEPC_{pro}$:mKOκ-NLS.* (m)-(o) $T_1$ plant expressing *$ZmPEPC_{pro}$:tdTomato-NLS*. (a), (d), (g), (j) and (m) show bright field. (b), (e), (h), (k) and (n) show each fluorescent protein (FP). (c), (f), (i), (l) and (o) show chlorophyll. Scale bars represent 10 μm.

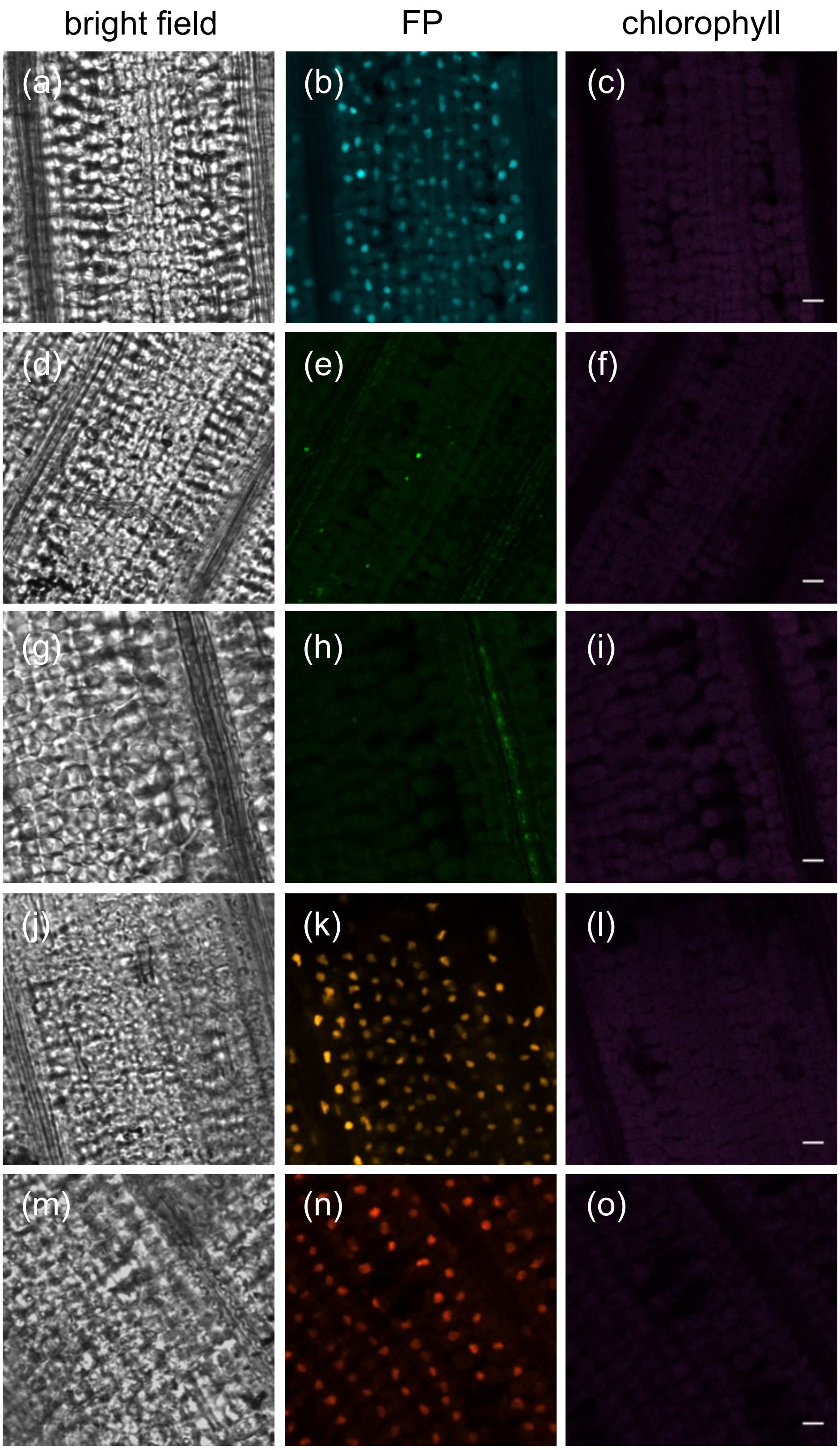


**Supplemental Figure 5: Fluorescent proteins expressed in mesophyll cells of ClearSee-treated leaves of stably transformed $T_1$ rice plants.** (a)-(c) Leaves expressing *pZmPEPC$_{pro}$::mTurquoise2-NLS.* (d)-(f) Leaves expressing *ZmPEPC$_{pro}$:mClover3-NLS.* (g)-(i) Leaves expressing *ZmPEPC$_{pro}$:mNeonGreen-NLS.* (j)-(l) Leaves expressing *ZmPEPC$_{pro}$:mKOκ-NLS.* (m)-(o) Leaves expressing *ZmPEPC$_{pro}$:tdTomato-NLS*. (a), (d), (g), (j) and (m) show bright field. (b), (e), (h), (k) and (n) show fluorescent protein (FP). (c), (f), (i), (l) and (o) show chlorophyll. Scale bars represent = 10 μm.

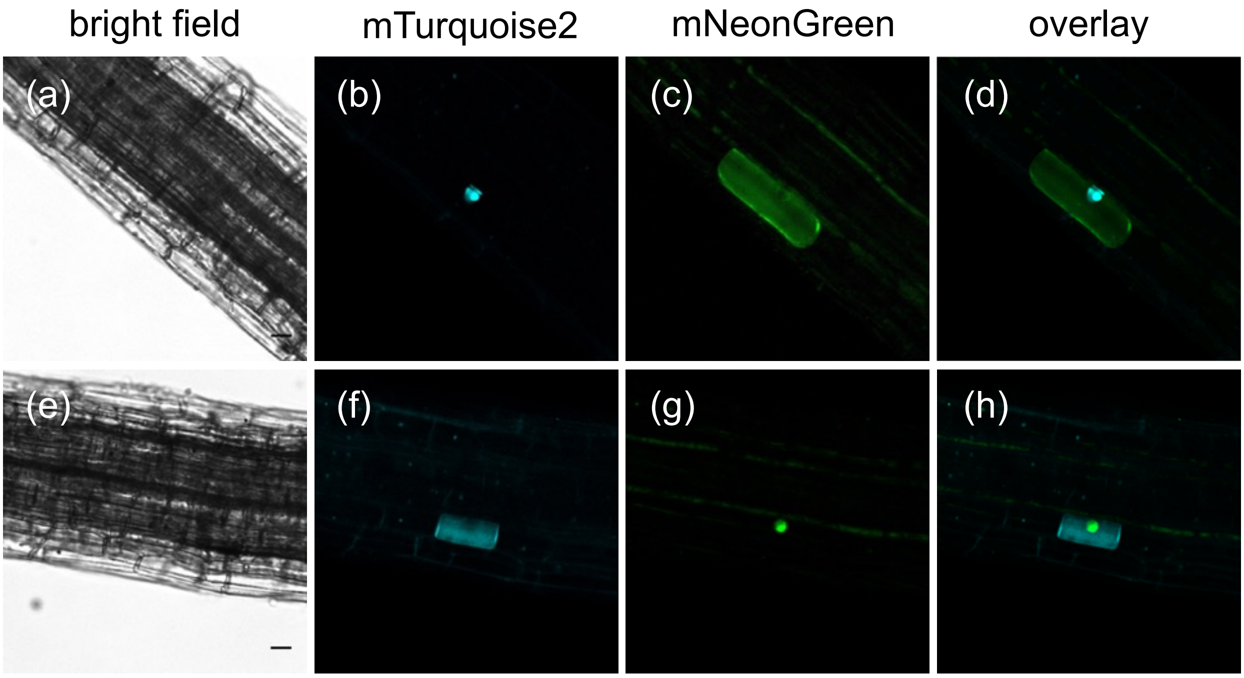


**Supplemental Figure 6: Fluorescent proteins targeted to different cell compartments in transiently transformed rice root cells.** (a)-(d) Root cell transformed with a construct expressing nuclear-localised mTurquoise2 and plasma membrane-localised mNeonGreen. (e)-(h) Root cell transformed with a construct expressing plasma membrane-localised mTurquoise2 and nuclear-localised mNeonGreen. Scale bars represent 10 µm.
