## Supplementary material for "Fluorescent reporters for functional analysis in rice leaves": S Table 1

**Table 1:** Screen of stably transformed $T_0$ plants expressing different fluorescent proteins. Numbers indicate independent $T_0$ plants for each fluorescent protein.

| Reporter | Nuclear signal | Autofluorescence only | Notes |
|---|---|---|---|
| mTurquoise2 | 5 strong / 3 weak | 1 | Clear nuclear signal, weak signal from autofluorescent structures |
| mNeonGreen | 5 strong / 4 weak | 0 | Clear nuclear signal, more signal from autofluorescent structures |
| mClover3 | 7 strong | 2 | |
| mKOκ | 7 strong | 0 | Clear nuclear signal, more signal from autofluorescent structures |
| tdTomato | 4 strong / 3 weak | 0 | |
| TagRFP-T | 3 strong / 3 weak | 0 | Non-robust nuclear signal |
| mCitrine | 2 weak | 3 | Weak nuclear signal for very few plants |
| mYPet | 1 weak | 4 | |
| mTFP1 | none detected | 6 | No nuclear signal detected |
| mRuby3 | none detected | 7 | |
| mKate2 | none detected | 7 | |
| mCardinal | none detected | 5 | |
